# The BRCA1-BARD1 complex associates with the synaptonemal complex and pro-crossover factors and influences RAD-51 dynamics during *Caenorhabditis elegans* meiosis

**DOI:** 10.1101/280214

**Authors:** Eva Janisiw, Maria Rosaria Dello Stritto, Verena Jantsch, Nicola Silva

## Abstract

During meiosis, the maternal and paternal homologous chromosomes must align along their entire length and recombine to achieve faithful segregation in the gametes. Meiotic recombination is accomplished through the formation of DNA double-strand breaks, a subset of which can mature into crossovers to link the parental homologous chromosomes and promote their segregation. Breast and ovarian cancer susceptibility protein BRCA1 and its heterodimeric partner BARD1 play a pivotal role in DNA repair in mitotic cells; however, their functions in gametogenesis are less well understood. Here we show that localization of BRC-1 and BRD-1 (*Caenorhabditis elegans* orthologues of BRCA1 and BARD1) is dynamic during meiotic prophase I; they ultimately becoming concentrated at regions surrounding the presumptive crossover sites, co-localizing with the pro-crossover factors COSA-1, MSH-5 and ZHP-3. The synaptonemal complex is essential for BRC-1 loading onto chromosomes but recombination is not. BRC-1 forms an in vivo complex with the synaptonemal complex component SYP-3 and the crossover-promoting factor MSH-5. Furthermore, BRC-1 is essential for efficient stage-specific recruitment of the RAD-51 recombinase to DNA damage sites when synapsis is impaired and upon induction of exogenous DNA double-strand breaks. Taken together, our data provide new insights into the localization and meiotic function of the BRC-1–BRD-1 complex and highlight their essential role in DNA double-strand break repair during gametogenesis.

**Author summary:** Sexually reproducing species rely on meiosis to transmit their genetic information across generations. Parental chromosomes (homologues) undergo many distinctive processes in their complex journey from attachment to segregation. The physiological induction of DNA double strand breaks is crucial for promoting correct chromosome segregation: they are needed to activate the DNA repair machinery responsible for creating physical connections, or crossovers (COs), between the homologues. In turn, crossovers promote the accurate segregation of the chromosomes in daughter cells. The BRCA1–BARD1 complex has a pivotal role during DNA repair in somatic cells and is exclusively located on unaligned chromosomal regions during mammalian meiosis. We show that in *Caenorhabditis elegans*, BRCA1 and BARD1 localize to chromosomes at all stages of meiotic prophase I and are enriched at presumptive crossover sites. We found that BRCA1 promotes DNA loading of the repair factor RAD-51 in specific mutant backgrounds and upon exogenous damage induction. Our data provide evidence for a direct physical association between BRCA1 and pro-crossover factors (including the synaptonemal complex) and identify an important role for BRCA1 in stimulating meiotic DNA repair. Further studies are necessary to identify the substrates acted upon by BRCA1–BARD1 complex to maintain genome stability in the gametes.

## Introduction

The genetic information encoded by DNA must be accurately copied and transmitted from one generation to the next. In somatic cells, DNA is duplicated and equally partitioned into daughter cells via mitosis, whereas in germ cells, which give rise to gametes, chromosome segregation relies on meiosis, a specialized cell division mechanism which produces haploid cells from diploid progenitors. Meiosis requires a unique programme of finely regulated events before cell division to accomplish faithful chromosome segregation. Cognate paternal and maternal chromosomes (homologous chromosomes) find each other (homologous pairing) and then fully align; the interaction is stabilized by formation of the synaptonemal complex (SC). Ultimately, exchange of DNA (recombination) between the homologues chromosomes establishes physical connections, which are essential for faithful segregation (1, 2).

The *Caenorhabditis elegans* gonad is a powerful system for studying chromosomes during both mitosis and meiosis because of the cytological accessibility and the spatio-temporal organization of nuclei into all prophase I stages (3). Morphological changes to chromosomes mark the engagement of key steps in meiotic progression. At meiotic onset, chromatin adopts a clustered, “half-moon” shape, reflecting chromosome movement and reorganization (4-6). This structure marks the transition zone (corresponding to the leptotene–zygotene stages). Once homologues are aligned, a tripartite proteinaceous structure called synaptonemal complex (SC) is formed between each homologue pair to allow genetic exchange during CO-dependent DNA repair (1, 2, 7-10). DNA recombination is initiated by the deliberate induction of DNA double-strand breaks (DSBs) by the topoisomerase II-like enzyme, SPO-11 (11, 12). In all species, the number of DSBs largely exceeds the final number of COs, suggesting that many DSBs are repaired via pathways such as inter-sister repair (IS) or synthesis-dependent strand annealing (13). In *C. elegans*, only one CO is formed between each homologous pair (14), and this depends on the function of the MSH-4/MSH-5 heterodimer (orthologues of the yeast and mammalian MutSγ complex components, MSH4/MSH5) (15-18), the cyclin COSA-1 (orthologue of mammalian CNTD1) (19, 20) and the E3 SUMO-ligase ZHP-3 (orthologue of yeast Zip3) (21-23). CO formation is abolished in absence of DSBs (e.g. in *spo-11* mutants) or synapsis; however, unlike in other model systems, lack of DNA breaks does not prevent SC formation in *C. elegans* (7, 11). Meiotic DSB repair also relies on RAD-51-mediated repair in *C. elegans* (24, 25): the RAD-51 recombinase localizes to discrete chromatin-associated foci starting in the transition zone and peaking in early pachytene; RAD-51 disengages from DNA in mid-pachytene (7). Markers of aberrant RAD-51 loading, such as increased foci number and/or extended accumulation, are bona fide indicators of defective DSB processing and recombination. CO induction triggers reorganization of the SC components into distinct domains on bivalents (pairs of homologous chromosomes held together by a chiasma): the central elements are confined to the short arm (containing the CO) and the axial elements to the long arm (26-30). This reorganization is particularly evident during diplotene, at which stage bivalents progressively condense and appear as six DAPI-stained bodies in diakinesis nuclei, which are a read-out for the successful execution of prophase I events (aberrant structures include achiasmatic chromosomes (univalents) or fused/fragmented chromatin masses (11, 16, 31)).

The breast and ovarian cancer susceptibility protein BRCA1 and its obligate heterodimeric partner BARD1 form an E3 ubiquitin ligase module (the BCD complex), the functions of which have been extensively studied in mitotic cells (32). The BRCA1–BARD1 heterodimer promotes homologous recombination (HR) during the S–G2 stages, by both favouring extended DNA break resection and preventing the non-homologous end joining (NHEJ)-promoting factor 53BP1 (33) from binding to the site of ongoing DNA repair. It also enhances BRCA2 and RAD51 loading at DNA damage sites to elicit accurate DNA repair (32). *BRCA1*-null mutants are embryonic lethal, thus hindering the study of this factor in gametogenesis (34-44). Mutants containing hypomorphic and gain-of-function alleles show increased apoptotic cell death during spermatogenesis, as well as reduced loading of the pro-CO factor MSH4 and a severe delay in MLH1 focus formation during oogenesis (45). *C. elegans brc-1/*BRCA1 mutants are viable and fertile, albeit with increased DNA damage-dependent apoptosis during oogenesis and SPO-11-dependent accumulation of RAD-51 foci, suggesting a defect in processing meiotic recombination intermediates (46, 47). Importantly, blocking *brc-1* function in CO-defective mutants leads to the formation of aberrant chromatin bodies in diakinesis nuclei, underscoring the importance of BRC-1 in the IS repair pathway (47).

Here we report that in the *C. elegans* germline, unlike in mammalian systems, BRC-1 and BRD-1 are abundantly expressed throughout meiotic prophase I and display a dynamic localization pattern in germ cells, switching from nucleoplasmic expression in early meiotic stages to SC association in pachytene, where they become progressively enriched at chromosome subdomains bearing the CO sites. We provide in vivo evidence that BRC-1 forms a complex with both MSH-5 and the SC central element, SYP-3. Localization of BRC-1 and BRD-1 in germ cells is differently regulated by synapsis and CO formation. Finally, we show that BRC-1 promotes stage-specific RAD-51 loading when SC formation is impaired and upon exogenous DNA damage induction. Similar findings are reported by Li and colleagues in the accompanying manuscript. Taken together, our data highlight the multiple functions of the BRC-1–BRD-1 heterodimer during gametogenesis.

## Results

### BRC-1 and BRD-1 display a dynamic localization pattern in the germline and are recruited to the short arm of the bivalent

To gain insight into BRC-1 and BRD-1 function during gametogenesis, we analysed their localization patterns during meiotic prophase I. To this end, we tagged the endogenous *brc-1* locus with a 3′ HA tag using a CRISPR/Cas9 approach (48, 49) and detected BRD-1 using a previously characterized specific antibody (46, 50). BRC-1::HA protein function was assessed by exposing the tagged animals to ionizing radiation (IR): as previously reported (46), *brc-1* mutants were sterile, whereas *brc-1*::*HA* worms responded to IR in a similar way to wild-type animals, thus proving that the tagged protein is fully functional (Fig 1A). Using a recently published method for isolating germline-enriched proteins (51) involving protein fractionation and western blot analysis, we showed that BRC-1::HA is enriched in the nucleus: most was in the soluble nuclear pool fraction and a smaller proportion was chromatin bound (Fig 1B). Interestingly, unlike in whole-cell extracts, BRC-1::HA was detected as a doublet in fractionated samples, suggesting that a less abundant isoform becomes detectable after enrichment with this extraction method or perhaps the presence of a post-translational modification.

**Fig 1.**
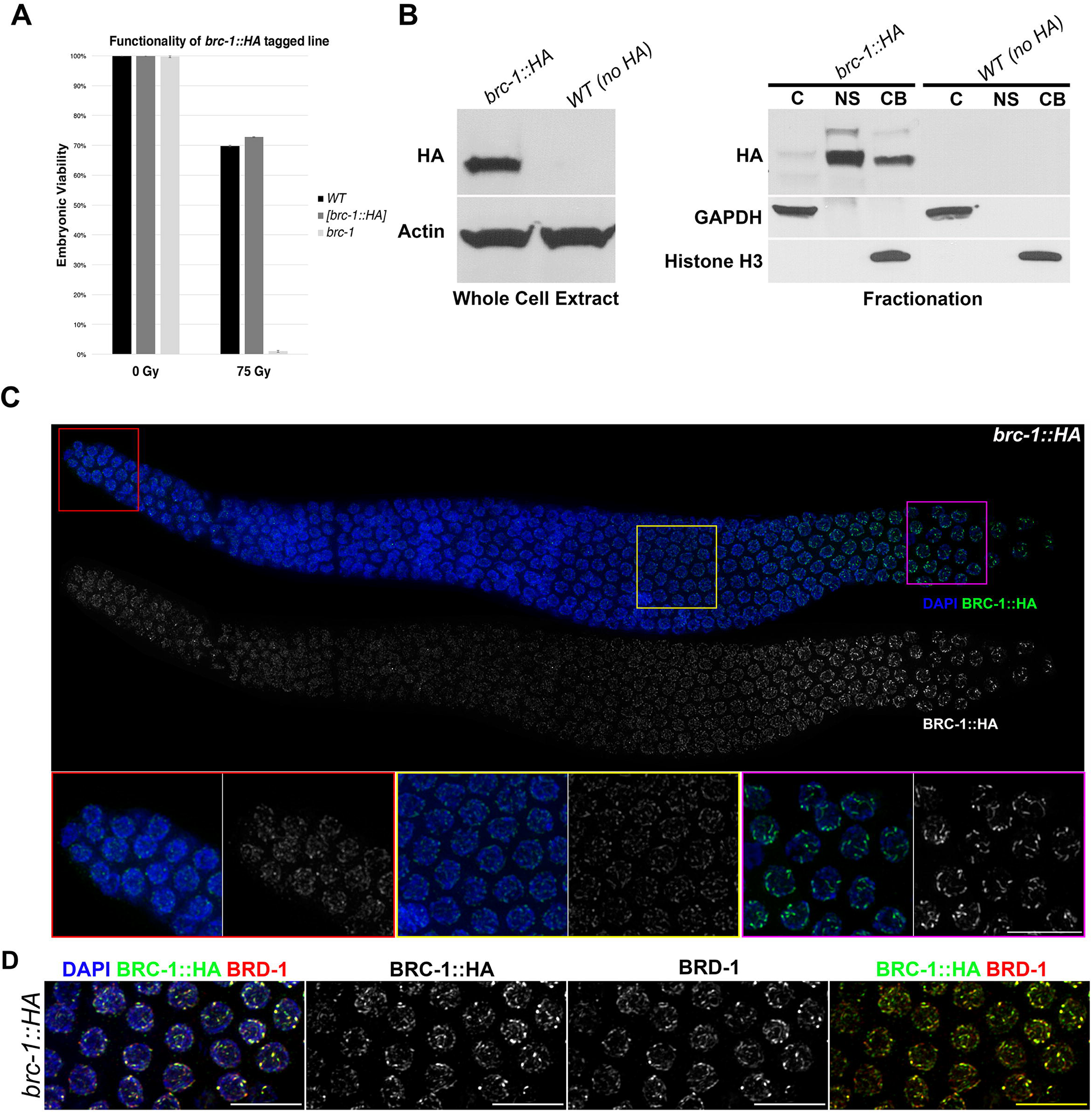
BRC-1–BRD-1 expression and localization during gametogenesis. (A) BRC-1::HA function was assessed by exposing worms to IR and then scoring the hatch rate of embryos laid in the following 24 hours. The mean of two independent experiments is shown. (B) Left: western blot analysis using an anti-HA antibody to monitor BRC-1::HA expression in whole-cell extracts. Actin was the loading control. Right: protein fractionation showing BRC-1::HA enrichment in the nucleus. Equal amounts of protein were loaded for each fraction. C = cytosol, NS = soluble nuclear pool, CB= chromatin-bound pool. GAPDH and histone H3 were used as loading controls for the cytosolic and chromatin-bound samples, respectively. (C) Top: whole-mount gonad from *brc-1*::*HA* worms dissected and stained with DAPI and anti-HA antibody, showing ubiquitous BRC-::HA expression throughout the germline. Note the progressive enrichment on the SC and short arms of bivalents. Bottom: enlarged images of specific regions of the gonad: mitotic tip (red frame), mid-pachytene (yellow frame) and late pachytene/diplotene (magenta frame). Scale bar, 10 μm. (D) Late pachytene nuclei stained with DAPI and anti-HA and BRD-1 antibodies display full co-localization of BRC-1::HA and BRD-1. Scale bar, 10 μm.

Previous reports indicate that during mouse meiosis, BRCA1 localizes to nascent SC elements during the leptotene/zygotene stages; in pachytene cells, it is exclusively located at asynapsed region of the XY-sex body during spermatogenesis or on asynapsed chromosomes during oocyte meiosis (52-54). In contrast, in the *C. elegans* germline both BRC-1 and BRD-1 were expressed in all nuclei during meiotic prophase I (Figs 1C and S1A) and, as expected, largely co-localized (Fig 1D). As observed in mammalian models, BRC-1 and BRD-1 loading is interdependent in nematodes: *brc-1* mutant germlines did not display any BRD-1 staining (Fig S1B) (55-57). Intriguingly, at the transition between mid to late pachytene, BRC-1 and BRD-1 staining switched from a rather diffuse to a discrete linear pattern along the chromosomes; in late pachytene nuclei, BRC-1 and BRD-1 progressively retracted into six short “comet-like” structures (Figs 1C, D and S1A), a specific pattern indicating localization to both CO sites and the short arm of bivalent (7, 8, 21, 22, 58, 59). To assess whether the BCD complex is indeed recruited to the short arm of the bivalent, we co-stained *brc-1*::*HA* germ lines with antibodies directed against the central element of the SC, SYP-1 (8) and the axial protein HTP-3 (60). As shown in Fig 2A, BRC-1 co-localized with SYP-1 in late pachytene/diplotene nuclei, confirming that the BCD complex becomes gradually concentrated in the region surrounding the CO site. Strikingly, BRC-1 enrichment at discrete regions preceded SYP-1 localization to the short arm of the bivalent (six robust stretches were seen only at late pachytene/diplotene stage). At meiosis onset, the PLK-2 polo-like kinase is enriched at the nuclear envelope attachment sites of chromosome ends, where it promotes homologous pairing and synapsis (61, 62). In late pachytene, PLK-2 re-locates to discrete domains along the SC, marking local enrichment of recombination factors (63). PLK-2 redistribution also occurs before SYP-1 redistribution to the short arm and influences the SC structure (63, 64). Given that BRC-1 redistribution had similar kinetics, we co-stained PLK-2 and BRC-1 (Fig 2B) and found that regions enriched for BRC-1 fully overlapped with the PLK-2 staining pattern in late pachytene and diplotene. Thus, the BCD complex is ubiquitously expressed during meiotic prophase I and becomes progressively enriched on the short arm of the bivalent prior to SYP-1 recruitment, where it co-localizes with PLK-2.

**Fig 2.**
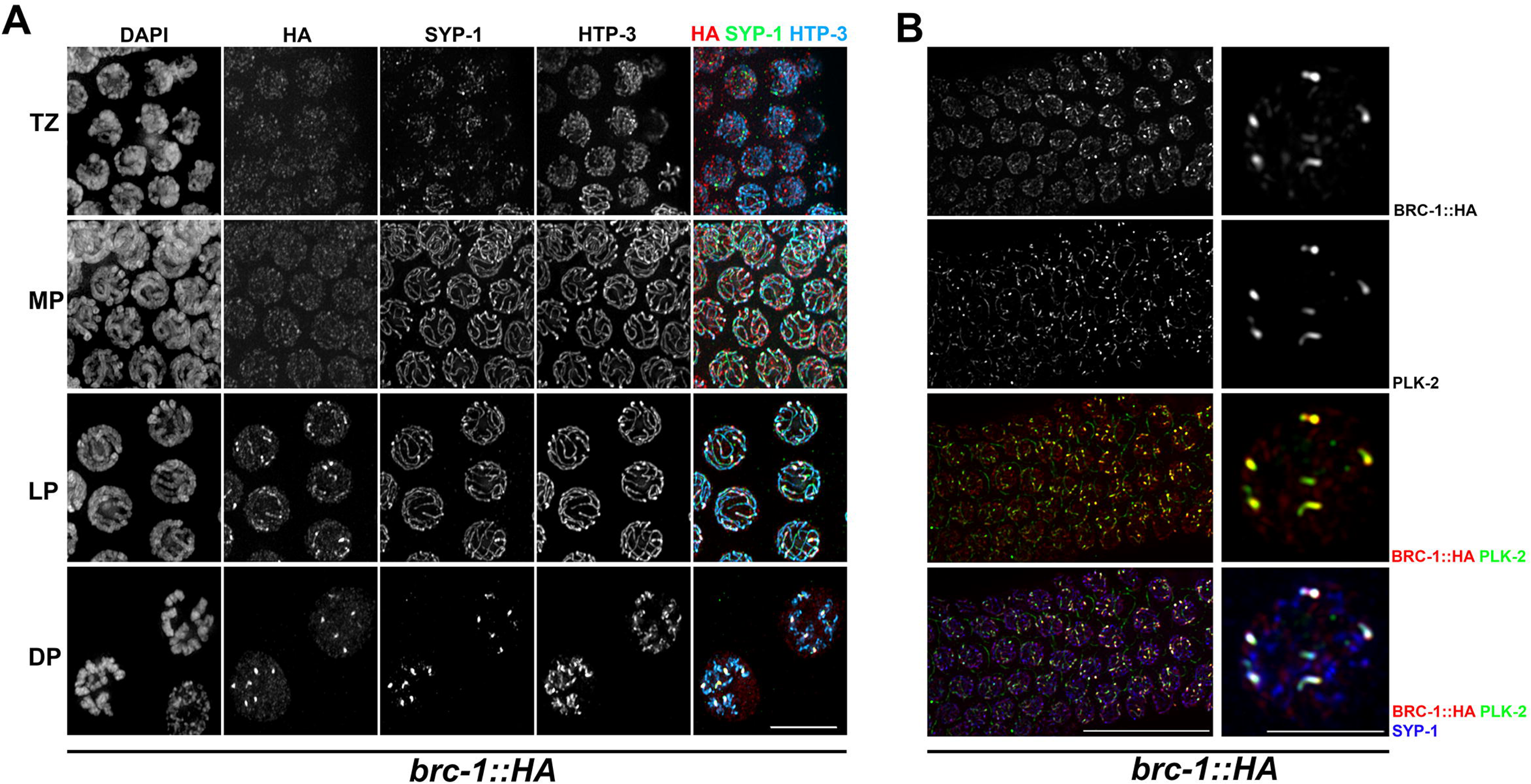
BRC-1 is enriched on the short arms of bivalents and co-localizes with SYP-1 and PLK-2. (A) BRC-1::HA, HTP-3 and SYP-1 localization patterns at different stages of meiotic prophase I. TZ = transition zone, MP = mid-pachytene, LP = late pachytene, DP = diplotene. Scale bar, 5 μm. (B) BRC-1::HA co-staining with anti-PLK-2 and anti-SYP-1 shows that BRC-1 is recruited concomitantly with PLK-2 and before SYP-1 to the shorts arm of bivalents. Scale bar, 30 μm (left) and 5 μm (right).

### CO establishment triggers redistribution of the BRC-1–BRD-1 complex

In *C. elegans*, formation of inter-homologue COs depends on several proteins, such as the COSA-1 cyclin (20), the MutSγ heterodimer, MSH4/MSH-5 (15, 16) and the ZHP-3 E3 SUMO-ligase (22). MSH-5 and ZHP-3 are detected at early meiotic stages, with the former accumulating in many foci (these are probably recombination intermediates with both CO and non-CO (NCO) outcomes) and the latter localizing along the SC (20-22). COSA-1 is prominently detected at mid–late pachytene transition as six foci (one CO for each homologue pair), which also contain MSH-5 and ZHP-3 (20). Since we observed BRC-1 and BRD-1 recruitment to the short arm of bivalents (chromosome subdomains caused by the formation of CO intermediates (26, 27, 29)), we wondered whether local enrichment of the BCD complex coincides with the regions labelled with pro-CO factors. Comparison of the localization dynamics of GFP::COSA-1 and BRC-1::HA showed that BRC-1 starts to become concentrated concomitantly with enhanced COSA-1 loading and defines a discrete area which later also contains SYP-1 (Fig 3A). We obtained the same localization pattern by monitoring BRD-1 loading (Fig S2). Furthermore, staining with anti-ZHP-3 antibody (21) also revealed full co-localization of ZHP-3 with BRC-1 (Fig 3A). To evaluate BRC-1 co-localization with MSH-5, we first added a 5′ GFP tag to the endogenous *msh-5* locus with CRISPR/Cas9. The tagged line was fully functional, with no defects in chiasmata formation (not shown), suggesting that GFP::MSH-5 is competent to promote CO formation. Similar to ZHP-3 and COSA-1, BRC-1::HA was enriched at defined regions containing a single GFP::MSH-5 focus, which also labels the CO site (Fig 3B). We performed structured illumination microscopy to further analyse BRC-1 association with the CO site. For this, we added a 5′ OLLAS tag to the endogenous *cosa-1* locus (65, 66). This fully functional line was crossed into *brc-1*::*HA* worms and co-stained for OLLAS (COSA-1), BRC-1 and SYP-1. This further confirmed BRC-1 enrichment around COSA-1-labelled CO sites; however, in these nuclei BRC-1 decorates the region of the SC embracing the putative recombination site; thus, it appears to surround, rather than overlapping with, COSA-1 (Fig 3C).

**Fig 3.**
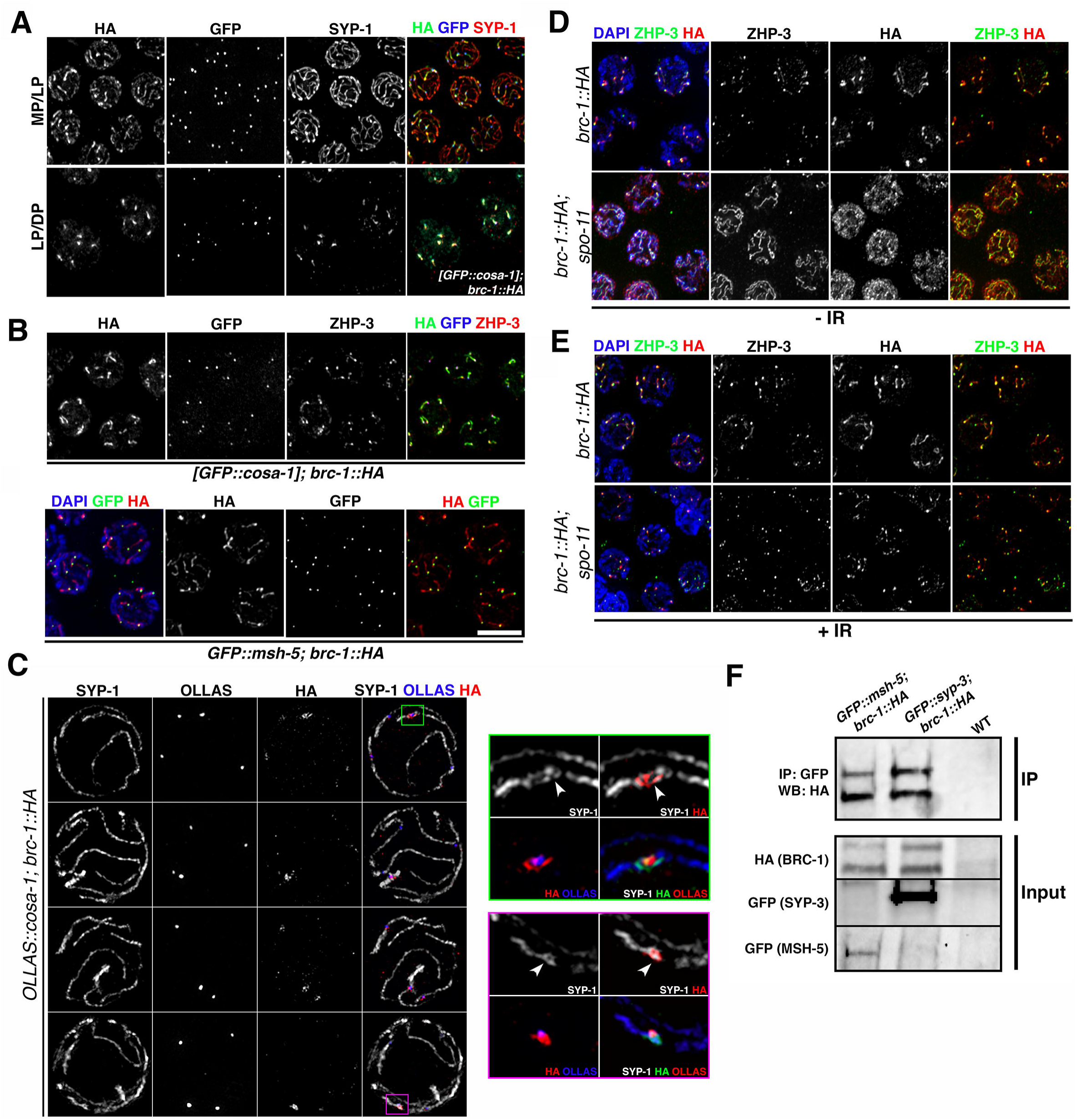
BRC-1 forms a complex with the CO machinery and the SC in vivo. (A) BRC-1::HA co-localizes with pro-CO factor COSA-1 in late prophase I. MP/LP = mid-/late pachytene, LP/DP = late pachytene/diplotene. Scale bar, 5 μm. (B) BRC-1::HA co-localizes with ZHP-3 and GFP::MSH-5 in late pachytene nuclei. Scale bar, 5 μm. (C) Partial projections of nuclei under super-resolution structured illumination microscopy: different examples show BRC-1::HA localization in the region surrounding the COSA-1-labelled CO site. BRC-1::HA forms a nodule-like structure together with SYP-1. (D) Late pachytene nuclei in non-irradiated samples show altered BRC-1::HA localization in *spo-11* mutants: BRC-1 and ZHP-3 remain localized along the SC without retraction to the short arms of bivalents. (E) Ionizing radiation rescues ZHP-3 and BRC-1::HA redistribution in *spo-11* mutants. (F) BRC-1::HA co-immunoprecipitates with GFP::MSH-5 and GFP::SYP-3 in vivo. GFP pull-downs were also performed in wild-type worms (WT) as a negative control.

To assess whether BRC-1–BRD-1 redistribution depends on CO establishment, we generated a *brc-1*::*HA*; *spo-11* mutant strain to monitor BRC-1::HA loading in absence of meiotic DSBs, which are essential for inducing CO formation. A previous report showed that in *spo-11* mutants COSA-1 occasionally forms very few foci (possibly arising from mitotic or spontaneous DSBs) and ZHP-3 remains localized along the SC without forming retraction “comets” due to a lack of chiasmata (20). In *spo-11* mutants, BRC-1 remained co-localized with ZHP-3 along the SC, without redistributing to chromosome subdomains. This confirms that BRC-1 redistribution depends on chiasmata formation (Fig 3D).

Exogenous DSB induction is sufficient to temporarily restore COSA-1 loading and therefore chiasmata formation in *spo-11* mutants (11, 20, 64). Thus, we investigated whether γ-irradiation could rescue the failure in BRC-1 redistribution. We exposed *brc-1*::*HA; spo-11* mutant worms to 20 Gy and analysed BRC-1 and ZHP-3 loading at 8 hours post irradiation: at this time point, all late pachytene nuclei in *spo-11* mutants display six COSA-1 foci, suggesting that CO induction is fully rescued (20). In the irradiated samples, ZHP-3 was retracted towards the CO site and, consistent with this, BRC-1 also became concentrated around the CO site (Fig 3E). Based on these data, we conclude that BRC-1 and BRD-1 localize to the short arms of bivalents and that their reorganization in mid-pachytene nuclei is dependent on CO establishment.

### BRC-1 physically interacts with MSH-5 and SYP-3 in vivo

Given its spatial association with both CO factors and the SC, we wondered whether BRC-1 formed protein complexes with these factors in vivo. We performed pull-down experiments using the *brc-1*::*HA; GFP*::*msh-5* strain (Fig 3) and crossed *brc-1*::*HA* into worms expressing a single-copy insertion transgene encoding a largely functional GFP::SYP-3 protein (67). Worms from both strains were used to generate cytosolic, soluble nuclear and chromatin-bound protein fractions (51): both nuclear fractions were pooled for immunoprecipitation experiments. Immunoprecipitation of GFP::MSH-5 and GFP::SYP-3 from *brc-1*::*HA; GFP*::*msh-5* and *syp-3; [GFP*::*syp-3]; brc-1*::*HA* strains, respectively, followed by western blot analysis with anti-HA antibodies revealed that that BRC-1::HA was present in both samples (Fig 3F). This suggests that BRC-1 forms a complex with both MSH-5 and SYP-3 proteins in vivo. Thus, we identified a previously unknown physical interaction of the BCD complex with the pro-CO factor MSH-5 and the SC central element SYP-3, highlighting a possible role for BRC-1–BRD-1 at the interface between synapsis and recombination.

### Synapsis and recombination have different effects on BRC-1 and BRD-1 loading

Given that CO establishment triggers BRC-1–BRD-1 redistribution (Fig 3C, D), we sought to analyse their localization in mutants that have impairment at different steps of CO formation. As already mentioned, an absence of DSBs leads to a lack of recombination, which prevents BRC-1 and BRD-1 retraction to the short arms of bivalents. We therefore asked whether impaired DNA repair by HR, but not by DSB induction, influences BRC-1 and BRD-1 localization. To address this, we crossed *brc-1*::*HA* into the *msh-5* mutant, which cannot convert recombination intermediates into mature CO products (7, 16). In *msh-5* mutants, BRC-1 accumulated along the SC but retraction was not observed (Fig 4A), similar to the localization pattern observed in *spo-11* (Fig 3). Then, we analysed BRC-1::HA staining in *rad-51* mutants, which have normal SC assembly but no homologous DNA repair due to lack of RAD-51-dependent strand displacement and invasion of the homologous chromosome (24, 25). Interestingly, BRC-1 had a rather punctate staining pattern, perhaps through labelling recombination-independent DNA joined molecules (Fig 4A). Despite this, a strong association with SYP-1 in chromosome subdomains was observed in nuclei exiting the pachytene stage (we also observed this in *msh-5* mutants). We observed a similar pattern of BRD-1 localization in *com-1* mutants (Fig S3): here, interfering with DSB resection impairs RAD-51 loading and therefore abolishes CO formation (68). These results suggest that a lack of COs per se impairs redistribution of the BCD complex in late pachytene cells without perturbing loading along the SC. However, in mutants such as *rad-51* that are defective in the early steps of recombination, BRC-1–BRD-1 association with the SC is also dramatically reduced. Next, we sought to analyse whether BRC-1 and BRD-1 loading is regulated by synapsis. We analysed BRC-1::HA loading in the complete and partial absence of SC, as well as in mutants in which synapsis occurs between non-homologous chromosomes. The central portion of the SC is formed by several proteins (SYP-1–4) which are loaded in an interdependent manner; thus, all are necessary to establish synapsis (7, 8, 58, 59). In the *syp-2* synapsis-null mutant (7), BRC-1::HA had a rather punctate staining pattern throughout meiotic prophase I. Strikingly, unlike in the wild type, where BRC-1 starts to spread along the SC immediately after the disappearance of RAD-51, in *syp-2* mutants BRC-1 foci remained in close proximity to and co-localized with RAD-51 foci in mid and late pachytene nuclei (Fig 4B). In *C. elegans*, a family of zinc-finger nuclear proteins connects chromosome-specific ends (i.e. pairing centres) to the nuclear envelope to promote chromosome pairing and synapsis (69, 70). ZIM-2 and HIM-8 bind to the ends of chromosomes V and X, respectively. Therefore, chromosome V is asynapsed in *zim-2* mutants and chromosome X is asynapsed in *him-8* mutants. We asked whether a partial deficiency in synapsis establishment (affecting only one chromosome pair) also changes BRC-1 loading dynamics. Analysis of BRC-1::HA expression in *him-8* and *zim-2* mutants revealed a lack of BRC-1 on unsynapsed chromosomes pairs, despite normal loading along the SC and retraction towards the CO site in the remaining bivalents (Fig 4C,D), suggesting that local synapsis defects do not impair BRC-1 loading. Lastly, we analysed BRD-1 loading in two mutants with deregulated SC assembly. HTP-1 is a HORMA-domain-containing protein essential to prevent SC assembly between non-homologous chromosomes and PROM-1 is an F-box protein involved in promoting meiotic entry and homologous pairing. Both *htp-1* and *prom-1* mutants display extensive SYP-1 loading between non-homologous chromosomes as well as asynapsed chromosome regions; consequentially, chiasmata formation is severely impaired (26, 71). Remarkably, the degree of BRD-1 co-localization with SYP-1 was extremely reduced in both *htp-1* and *prom-1* mutants, with most BRD-1 detected as bright agglomerates within the nucleus (Fig S4). Thus, we conclude that BRC-1 and BRD-1 redistribution during meiotic progression requires CO establishment and is tightly regulated by SCs.

**Fig 4.**
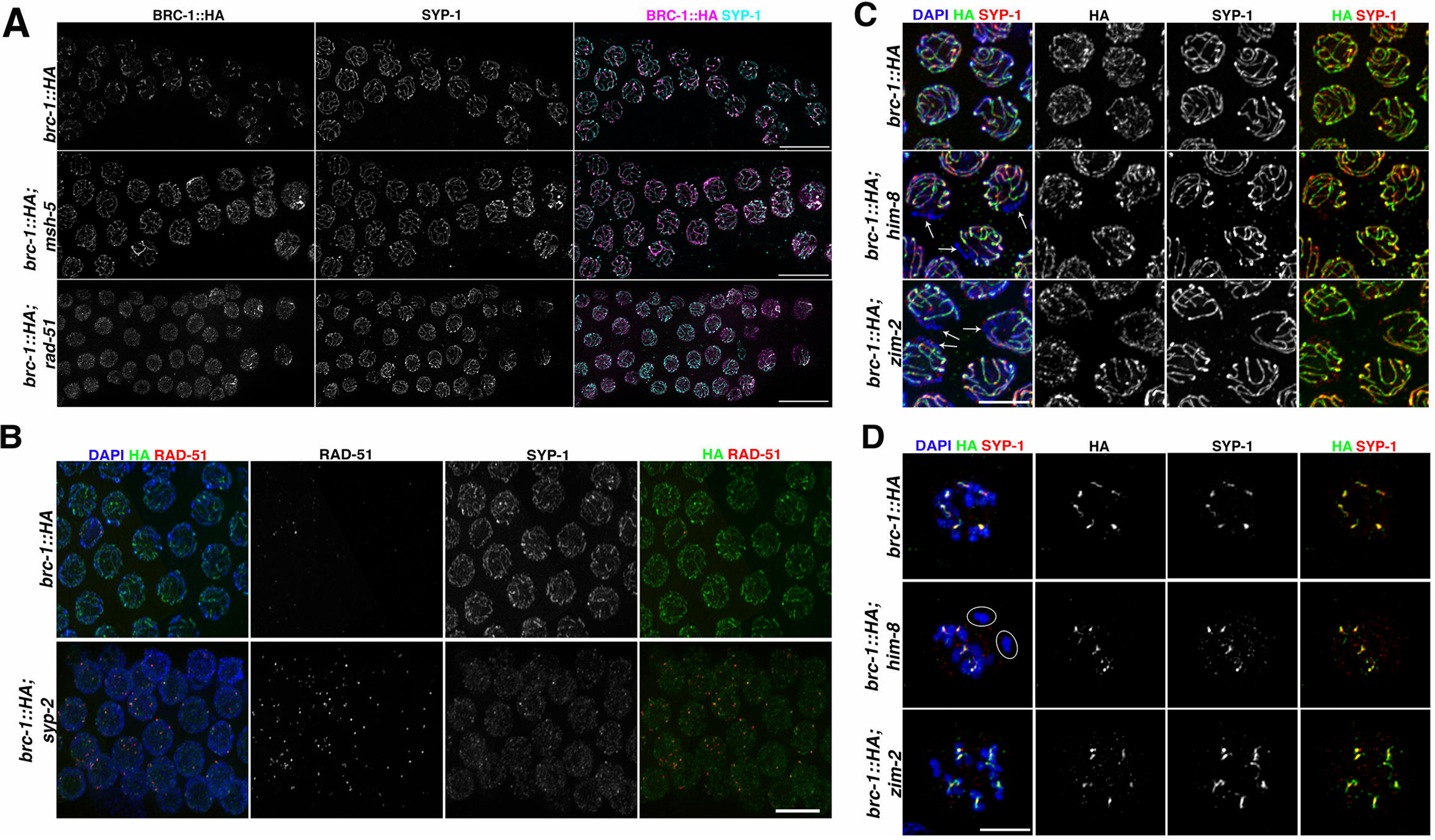
Recombination and synapsis differentially regulate BRC-1 localization. (A) BRC-1::HA localization was assessed in *msh-5* and *rad-51* mutants. In the *msh-5* mutant, BRC-1::HA accumulates in the SC, as in *spo-11* mutants; in the *rad-51* mutant, it displays interspersed staining, and association with the SC is strongly reduced. Scale bar, 10 μm. (B) Abrogation of synapsis triggers BRC-1::HA accumulation into discrete chromatin-associated foci which co-localize with RAD-51 in late pachytene cells. Scale bar, 5 μm. (C) Unlike in the *him-8* or *zim-2* mutant, respectively, BRC-1::HA does not accumulate on asynapsed chromosome X or V in late pachytene nuclei. Arrows indicate regions of DNA devoid of both SYP-1 and BRC-1::HA. Scale bar, 5 μm. (D) Diplotene nuclei of *him-8* and *zim-2* mutants clearly lack BRC-1::HA on asynapsed univalents (circled). Scale bar, 5 μm.

### BRC-1 promotes RAD-51 recruitment in the absence of synapsis

BRC-1 is dispensable for establishing synapsis and chiasmata; however, *brc-1* mutant germlines have a higher number of and more persistent RAD-51-labelled recombination intermediates compared with the wild type (Fig S5) (46, 47). Impaired BRC-1 localization, and probably also impaired function, in CO-defective mutants leads to the formation of abnormal chromosome structures in diakinesis nuclei, possibly due to deficient IS repair (47). DSB repair during meiosis is channelled into both CO and NCO pathways. Since it has been suggested that BRC-1 might preferentially function in NCOs (47), we investigated whether other factors involved in resolving the recombination intermediates required for both CO and NCO repair might also be affected. In somatic cells, the RTR complex mediates efficient resolution of recombination intermediates by promoting the dissolution of double Holliday junctions to yield non-CO products (72-74). RMI1 is an essential component of the RTR complex and a scaffolding component for other complex members, BLM and TOP3A, which promotes their dissolution activity (74). The *C. elegans* RMI1orthologue, RMH-1, localizes to recombination foci during meiosis: it appears in early pachytene and peaks in mid-pachytene, accumulating in many foci and possibly labelling all recombination intermediates. At late pachytene transition, the number of RMH-1 foci is reduced to roughly six per nucleus; these foci co-localize with foci of the pro-CO factors COSA-1, MSH-5 and ZHP-3. Lack of RMH-1 causes a drastic reduction in chiasmata formation due to impaired COSA-1 and MSH-5 loading. However, in CO-deficient backgrounds such as *cosa-1, msh-5* and *zhp-3* mutants, RMH-1 is still recruited in early pachytene but is not retained until late pachytene. Therefore, it has been postulated that RMH-1 functions in both the CO and NCO pathways (75). MSH-5 displays a similar localization, but does not fully co-localize with RMH-1 (20, 75). We scored COSA-1, MSH-5 and RMH-1 nuclear localization in *brc-1* mutants in nuclei spanning the transition zone to late pachytene stage. Interestingly, GFP::MSH-5 accumulation was reduced in early and mid-pachytene, with a similar, but less prominent, trend for GFP::RMH-1 (Fig 5A,B). By late pachytene, both proteins had been recruited into six foci, together with COSA-1, suggesting that the early processing of recombination intermediates might be defective in absence of BRC-1.

**Fig 5.**
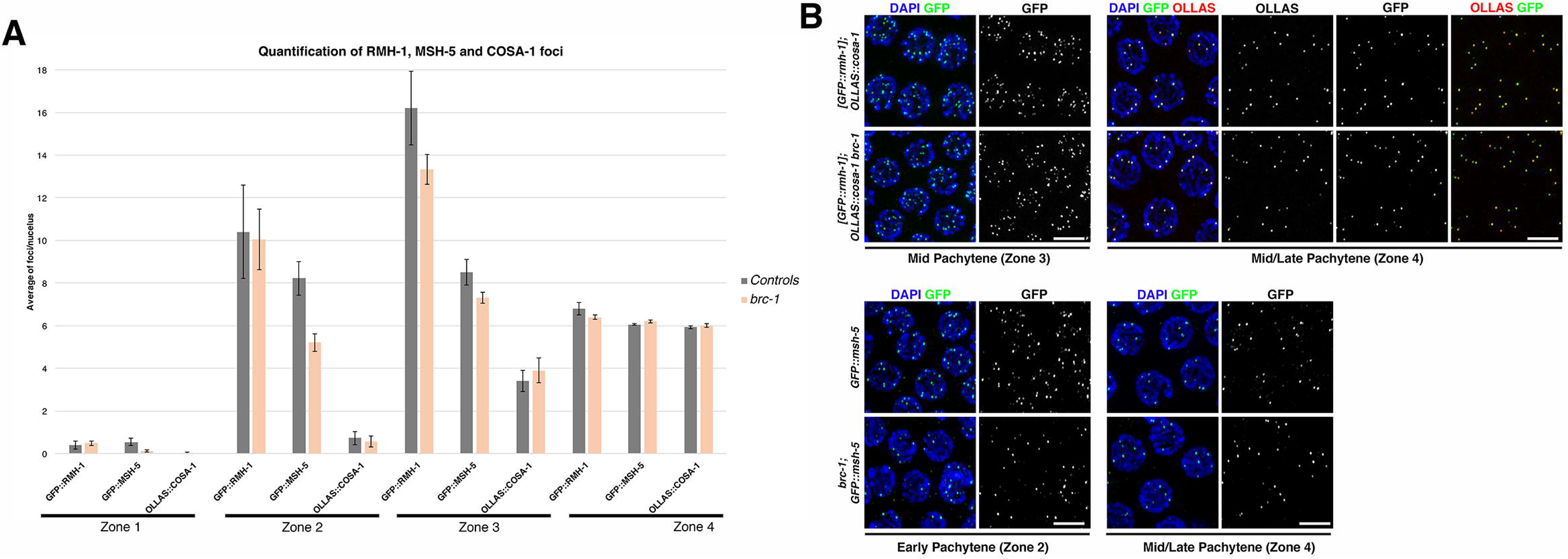
Analysis of recombination markers in *brc-1* mutants. (A) Quantification of GFP::RMH1, GFP::MSH-5 and OLLAS::COSA-1 markers in *brc-1* mutants and control animals. Gonads were divided into four equal regions from the transition zone to the late pachytene stage. The average number of foci per nucleus from at least three gonads per genotype is shown. For GFP::RMH-1 and OLLAS::COSA-1 quantification, the number of nuclei scored for each gonad region in the controls (and *brc-1* mutants) were: zone 1, 242 (334); zone 2, 185 (244); zone 3, (181 (214); zone 4, 124 (136). For GFP::MSH-5 quantification, the equivalent numbers were: zone 1, 230 (403); zone 2, 210 (410); zone 3,165 (303); zone 4, 121 (147). Error bars show S.E.M. (B) Representative nuclei at different meiotic stages co-stained for GFP::RMH-1 with OLLAS::COSA-1 (upper panels) or GFP::MSH-5 (lower panels). Scale bar, 5 μm. Note that both GFP::RMH-1 and GFP::MSH-5 are expressed in fewer foci in early and mid-pachytene but not in late pachytene in *brc-1* mutants.

Given that BRC-1 and BRD-1 loading are regulated by synapsis and the establishment of COs, and that a lack of BRC-1 might affect the processing of NCOs rather than COs, we next assessed the effects of BRC-1 depletion in genetic backgrounds defective in chiasmata formation, which hence rely solely on NCOs to repair meiotic DSBs. We first analysed DAPI-stained bodies in diakinesis nuclei from *cosa-1 brc-1* and *brc-1; syp-2* double mutants to confirm the presence of aberrant chromatin structures (Fig 6A), as previously reported (46, 47). As abnormalities in diakinesis nuclei can result from impaired RAD-51-dependent repair of meiotic DSBs (24, 31, 76), we sought to analyse whether lack of *brc-1* altered RAD-51 dynamics. To this end, we quantified RAD-51 in *cosa-1 brc-1* and *brc-1; syp-2* mutants. Failure to convert recombination intermediates into mature CO products has been linked to increased RAD-51 levels and its delayed removal during meiotic prophase due to either excessive DSB induction or slower processing of recombination intermediates (5, 6, 15, 16), which are eventually channelled into alternative repair pathways (e.g. IS repair) (7). In fact, both *cosa-1* and *syp-2* mutants accumulated high levels of RAD-51, which disengaged from chromatin in mid and late pachytene, respectively (Fig 6B, C) (7, 20). Remarkably, removal of BRC-1 from *cosa-1* and *syp-2* mutants had different effects on RAD-51 dynamics: in both *cosa-1 brc-1* and *brc-1; syp-2* double mutants, there were far fewer RAD-51 foci in early pachytene compared with both single mutants; however, in *cosa-1 brc-1* mutants RAD-51 accumulation was dramatically prolonged until diplotene, whereas in the *brc-1; syp-2* mutant overall RAD-51 staining was dramatically reduced (Fig 6B, C). Aberrant chromosome structures occurred at a particularly high frequency in *brc-1; syp-2* mutants, consistent with the severe reduction in RAD-51 loading in pachytene nuclei (Fig 6A). Thus, in CO-defective mutants, BRC-1 regulation of RAD-51 dynamics is altered by the presence of the SC.

**Fig 6.**
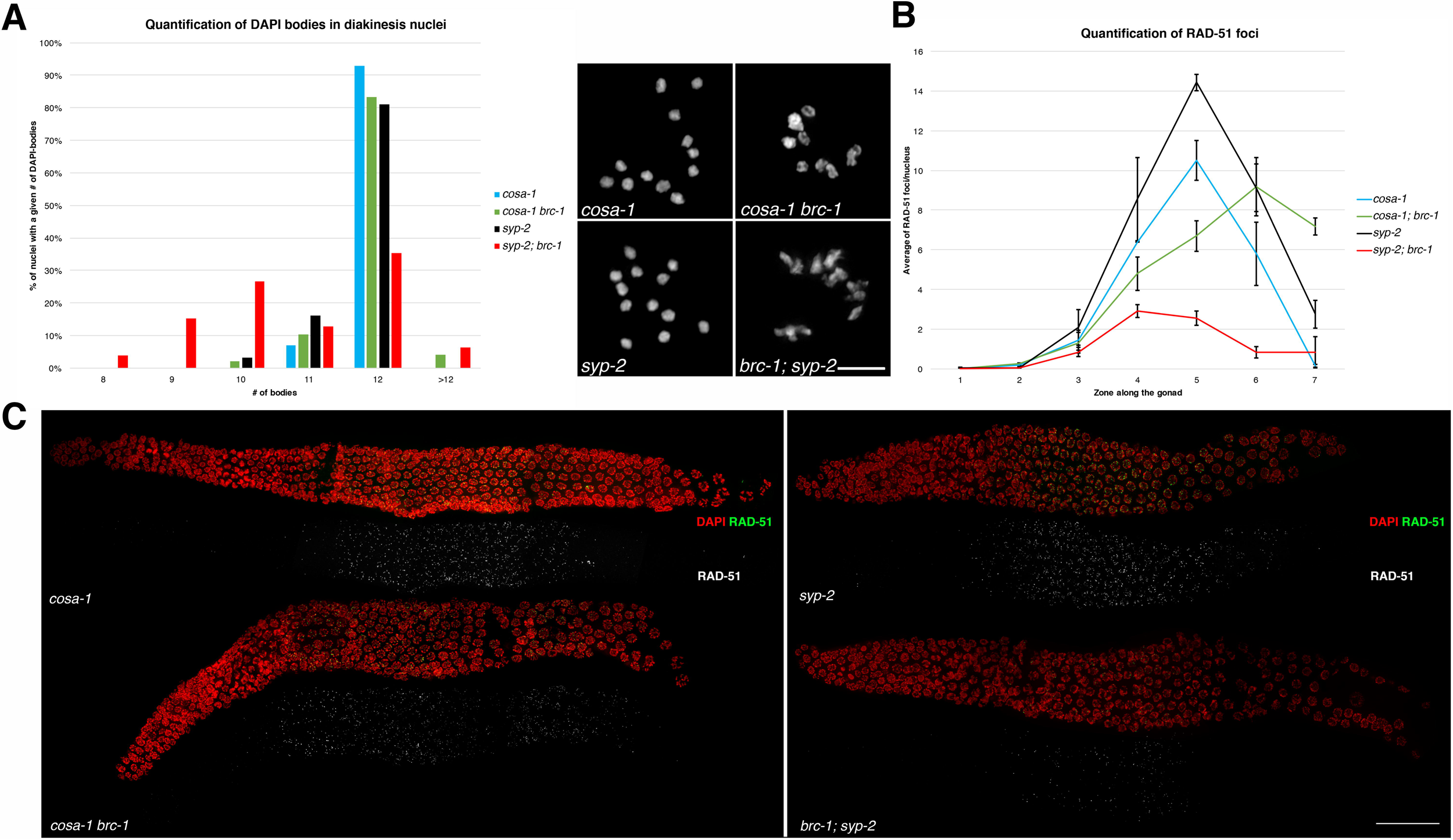
Loss of BRC-1 differently influences RAD-51 loading in *cosa-1* and *syp-2* mutants. (A) Left: quantification of DAPI-stained bodies in diakinesis nuclei in different genotypes. Number of diakinesis nuclei scored: *cosa-1*, 42; *cosa-1 brc-1*, 48; *syp-2*, 31; *brc-1; syp-2*, 79. Right: representative images of DAPI-stained diakinesis nuclei. Scale bar, 3 μm. (B) Quantification of RAD-51 foci per nucleus throughout the germline. Each gonad was divided into seven equal zones and RAD-51 foci were counted in each nucleus. Data show the average of at least three gonads for each genotype. Number of nuclei scored from zone 1 to zone 7 in different mutants: *cosa-1* – 162, 225, 179, 165, 142, 107, 124; *cosa-1 brc-1* – 347, 335, 285, 274, 228, 175, 149; *syp-2* – 244, 265, 241, 233, 190, 131, 118; *brc-1; syp-2* – 226, 305, 275, 297, 254, 161, 147. Error bars show S.E.M. (C) Whole-mount gonad stained with DAPI and anti-RAD-51 antibody. Note accumulation of RAD-51 foci in late pachytene in *cosa-1 brc-1* double mutants, which is not observed in *cosa-1* single mutants. In *brc-1; syp-2* animals, the number of RAD-51 foci was dramatically reduced. Scale bars, 30 μm.

### Efficient RAD-51-mediated repair upon exogenous DSB induction requires functional BRC-1

Exposure of *brc-1* and *brd-1* mutants to IR causes dose-dependent hypersensitivity which eventually culminates in full sterility, possibly due to the formation of highly unstructured chromatin bodies in diakinesis nuclei (46). These structures resemble those formed upon BRC-2/BRCA2 depletion, which in worms is essential for RAD-51 loading (31, 76), and COM-1/Sae2 depletion, which promotes DSB resection (68, 77). Both mutants lack RAD-51 recruitment onto DNA during meiotic prophase I. We therefore sought to investigate whether the aberrant chromatin masses observed in irradiated *brc-1* mutants were caused by impaired RAD-51 recruitment. To be efficiently loaded to the single-stranded DNA (ssDNA) tails generated after resection, RAD-51 must be exchanged with RPA (RPA-1 in worms), which coats ssDNA tails to stabilize them and prevent DNA from self-winding (78, 79). We generated a *brc-1* mutant strain expressing RPA-1::YFP (80) and analysed RAD-51 and RPA-1 loading at two different time points post irradiation. We observed a dramatic reduction in RAD-51 focus formation specifically in mid to late pachytene nuclei of *brc-1* mutants, along with enhanced RPA-1 levels (Fig 7A). At 24 hours post irradiation, both RAD-51 and RPA-1 were still abundant in *[rpa-1*::*YFP]* animals; in contrast, in *brc-1; [rpa-1*::*YFP]* mutants RPA-1 was still expressed at higher levels than in controls, but RAD-51 was remarkably reduced (Fig 7B). Prompted by these results, we decided to analyse the loading dynamics of BRC-1::HA and RAD-51 after IR exposure to assess whether exogenous DSB formation affects the mutual spatio-temporal regulation of these proteins. Under physiological growth conditions, BRC-1 and RAD-51 localization did not overlap prior to BRC-1 enrichment in the SC, which occurs after RAD-51 disappearance (Fig S6A, B). At 1 hour post irradiation, BRC-1::HA started to form discrete chromatin-associated foci in pre-meiotic nuclei, often in close proximity to (but not co-localizing with) RAD-51 foci (Fig S6A,B). Although abundant RAD-51 accumulation was triggered by IR exposure throughout the germline, BRC-1::HA levels were only modestly increased. However, western blot analysis revealed a shift in BRC-1::HA migration after IR which remained unchanged throughout the time course (Fig S6C), suggesting that exogenous DNA damage might elicit post-translational modifications of BRC-1. Western blot analysis also showed a slight increase in BRC-1::HA abundance, confirming our immunofluorescence data (Fig S6A). In meiotic nuclei, BRC-1 was detected along the SC at an earlier time point than in non-irradiated animals, but retraction towards the short arms of bivalents appeared delayed (Fig S6A). Samples analysed 8 hours after IR revealed robust BRC-1 and RAD-51 co-localization in nuclei residing in the mitotic tip; however, as at the earlier time point, no clear co-localization was observed in pachytene nuclei (Fig S6A, B). At 24 hours post irradiation, BRC-1::HA foci in the mitotic nuclei had largely disappeared and bright RAD-51 foci were observed only in enlarged, G2-arrested nuclei that were still undergoing repair; in contrast, bright RAD-51 foci co-localizing with BRC-1 were occasionally seen in non-arrested nuclei. Nuclei progressing through meiotic prophase I displayed more BRC-1 accumulation along chromosomes at 24 hours post irradiation compared with unirradiated controls. Taken together, our observations revealed that BRC-1 accumulation in the germline is modulated by exogenous DNA damage and that the clear BRC-1 and RAD-51 co-localization observed only in mitotic nuclei was cell cycle dependent.

**Fig 7.**
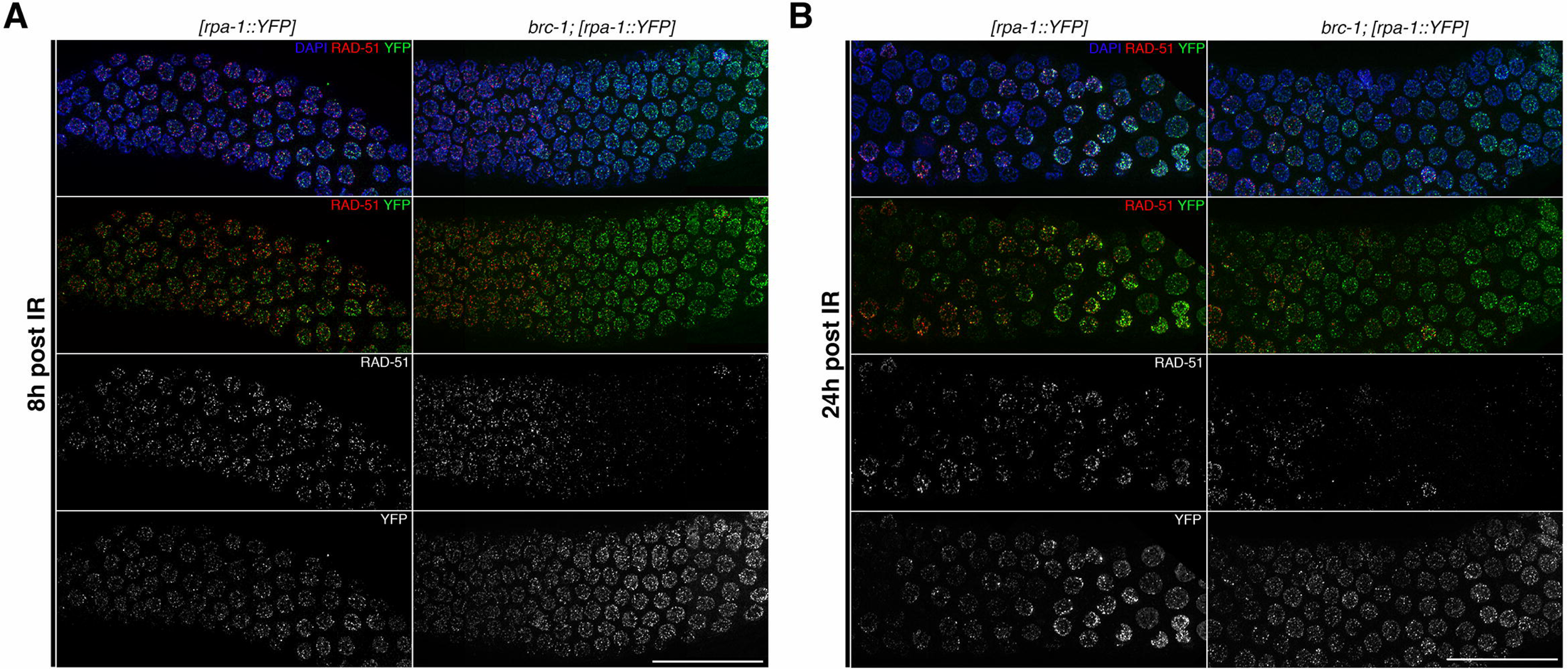
Efficient accumulation/exchange of RAD-51 and RPA-1 upon exogenous DNA damage requires BRC-1 function. (A) Time course analysis of RAD-51 and RPA-1::YFP DNA loading in irradiated *brc-1* and controls. Worms were irradiated with 75 Gy IR and analysed after 8 hours. Nuclei in mid-pachytene display enhanced RPA-1 levels and drastically reduced RAD-51 levels in *brc-1* mutants compared with controls. (B) The same analysis performed at 24 hours post irradiation showed severely reduced RAD-51, with higher RPA-1 levels that were identical to those at the earlier time point. Scale bars, 30 μm.

## Discussion

Our study sheds new light on the expression dynamics of the *C. elegans* BRC-1–BRD-1 heterodimer during meiotic prophase I and reveals that BRC-1 regulates RAD-51 accumulation in the germline under both unchallenged conditions and upon exogenous DNA damage induction. We show that in contrast to mammalian meiosis, where BRCA1 is loaded exclusively at asynapsed chromosome regions during spermatogenesis and oogenesis in pachytene cells (52-54), in worms both BRC-1 and BRD-1 are expressed throughout meiotic prophase I and are progressively enriched on the short arms of bivalents, in a CO- and SC-dependent manner. Our data provide the first evidence that BRC-1 forms a complex in vivo with the pro-CO factor MSH-5 and the SC central element SYP-3. Taken together, our findings provide new insight into the meiotic functions of BRC-1 and BRD-1 and show that the BCD complex is essential for preserving genome integrity and stimulating HR during gametogenesis.

### The BCD complex functions at the interface of synapsis and recombination

BRC-1 and BRD-1 display a highly dynamic localization pattern during meiotic prophase I progression, shifting from a pattern of rather diffuse accumulation at early stages to a robust association with the SC, which culminates in retention of the BCD complex at the region of the bivalent harbouring the chiasma (Figs 1–3). Remarkably, accumulation of BRC-1–BRD-1 at specific chromosomal subdomains occurred prior to retraction of the SC central elements to those domains but was concomitant with recombination factor-dependent enrichment of PLK-2 at the SC (Fig 2B) (63, 64), suggesting that the BCD complex is actively targeted to the region surrounding the CO rather than passively recruited following SC remodelling. The fact that recruitment of BRC-1–BRD-1 to the region surrounding the chiasma has similar kinetics to PLK-2 recruitment and precedes SYP-1 redistribution suggests that the BCD complex (i) is brought into place via physical interaction with the CO machinery (Fig 3); (ii) might respond to changes in the physical properties of the SC, as triggered by chiasmata formation or (iii) might be directly induced by PLK-2.

Importantly, Li et al. (accompanying manuscript) observe the same localization pattern for BRC-1 and BRD-1 by employing GFP-tagged functional lines. However, in contrast with the aforementioned study in which they analysed localization in live worms, we did not detect BRD-1 loading in fixed *brc-1* mutant germlines, confirming what was previously reported in (50). This difference might be possibly due to distinctive processing of the samples or tag-dependent alterations of the properties of the fusion proteins (81).

Our data favour a model in which the SC is essential for initial recruitment of the BCD complex onto the chromosomes and later accumulates at the CO site due to the local concentration of recombination factors. In fact, BRC-1 recruitment to the SC is not prevented in *msh-5* or *spo-11* mutants (both of which are defective in CO formation but proficient in synapsis establishment). However, similar to ZHP-3, BRC-1 fails to retract (Figs 3C and 4A). Irradiation of *spo-11* mutants restored BRC-1 and ZHP-3 redistribution to the short arms of bivalents (Fig 3D), confirming that CO establishment per se is the key trigger of local BCD complex enrichment. Abrogation of synapsis dramatically changed the BRC-1 expression pattern: it remained punctate throughout meiotic prophase I and displayed extensive and specifically co-localization with RAD-51 in late pachytene cells (Fig 4B). However, in mutants in which only one chromosome pair was asynapsed, such as *him-8* and *zim-2* mutants, BRC-1 was not loaded onto the unsynapsed regions but loading dynamics were normal for the other chromosome regions (Fig 4C, D). It was recently shown that PLK-2 plays a pivotal role in modulating the physical state of the SC in response to recombination and that an absence of synapsis impairs PLK-2 redistribution from the nuclear envelope to chromosome subdomains (63, 64, 82), which might explain the different BRC-1 localization patterns in *syp-2* mutants. Different BRD-1 localization patterns were observed in *htp-1* and *prom-1* mutants, but both were characterized by extensive non-homologous synapsis. BRD-1 accumulated in bright agglomerates in the nucleus, suggesting that SYP loading per se is not sufficient to recruit BRC-1–BRD-1 onto the SC (Fig S6).

### Crosstalk between the BCD complex and RAD-51 is governed by the SC

Blocking BRC-1 function had opposing effects on the progression of recombination intermediates in *cosa-1* and *syp-2* (CO-defective) mutants. RAD-51 accumulation was exacerbated in *cosa-1* single mutants and largely suppressed in *syp-2* mutants (Fig 6), leading to the formation of aberrant chromatin masses in diakinesis nuclei both mutant backgrounds, as previously reported (47). Based on genetic data, BRC-1 function was previously postulated to be essential for IS repair of meiotic DSBs (47, 83, 84); our data corroborate this model. In *cosa-1 brc-1* double mutants, the presence of an intact SC might still impose a homologue-biased constraint for an inter-homologue, CO-independent pathway that relies on RAD-51-mediated repair but not on BRC-1 function. However, in the absence of synapsis, repair of recombination intermediates is probably channelled entirely through the IS repair pathway because the sister chromatid is the only available repair template: SC depletion triggers association of BRC-1 with RAD-51 in late pachytene cells at presumptive repair sites, thereby promoting HR-mediated repair. We also observed fewer RAD-51 foci in *brc-1; syp-2* double mutants during early pachytene, suggesting that BRC-1 is nonetheless required to (directly or indirectly) promote efficient RAD-51 loading, although co-localization with RAD-51 at meiotic onset might be very transient. We observed that a lack of BRC-1 reduces the loading of recombination markers such as MSH-5 and RMH-1 in early pachytene, suggesting that even in the presence of the SC, BRC-1–BRD-1 function is required to efficiently promote the processing of recombination intermediates. Moreover, in *brc-1* mutants exposed to exogenous DSB induction, RAD-51 is not efficiently retained in mid- to late pachytene cells (Fig 7). This is not due to impaired resection, as shown by the abundant recruitment of RPA-1, which stabilizes ssDNA. However, RAD-51 loading is comparable to controls in later stages, suggesting that stabilization, rather than loading per se, might require the action of the BCD complex. This is in line with the findings reported by Li et al. (see accompanying manuscript).

When we scored BRC-1 levels after exposure to IR, we detected a slight increase in abundance but a marked difference in protein migration on western blots (Fig S6), suggesting that exogenous DNA damage promotes post-translational modification of BRC-1. Importantly, despite dramatically enhanced RAD-51 levels upon irradiation, we observed clear co-localization with BRC-1 only in mitotic cells and not during pachytene, once again confirming that these proteins co-localize only when the SC is indeed absent (Fig S6). Our findings suggest that the BCD complex responds to both synapsis and recombination and that the SC might act as a docking site for the BRC-1–BRD-1 complex to modulate its function in promoting DNA repair.

## Materials and methods

### Worm strains

All the worm strains used were grown at 20°C and the N2 Bristol strain was used as the wild type. The following alleles were used: LGI: *syp-3(ok758), prom-1(ok1140)*; LGII: *[GFP*::*rmh-1]* (75), *[GFP*::*syp-3]* (67), *[GFP*::*cosa-1]* (20); LGIII: *brc-1(tm1145), brc-1*::*HA* (this study), *brd-1(gk297), cosa-1(tm3298), OLLAS*::*cosa-1* (this study), *com-1(t1626);* LGIV: *spo-11(ok79), him-8(tm611), zim-2(tm574), msh-5(me23), GFP*::*msh-5* (this study), *htp-1(gk174), rad-51(lg8701)*; LGV: *syp-2(ok307)*. No information is available on the chromosomal integration of *[rpa-1*::*YFP]* (80).

### Cytological procedures

For cytological analysis of whole-mount gonads, age-matched worms (24 hours post-L4 stage) were dissected in 1× PBS on a Superfrost Plus charged slide and fixed with an equal volume of 2% PFA in 1× PBS for 5 min at room temperature. Slides were freeze-cracked in liquid nitrogen and then incubated in methanol -20°C for 5 min, followed by three washes in PBST (1× PBS, 0.1% Tween) at room temperature. Slides were blocked for 1 hour at room temperature in PBST containing 1% BSA and then primary antibodies were added in PBST and incubated overnight at 4°C. Slides were then washed in PBST at room temperature and secondary antibodies were applied for 2 hours. After three washes in PBST for 10 min each, 60 μl of a 2 μg/ml stock solution of DAPI in water was added to each slide and stained for 1 min at room temperature. Samples were washed again for at least 20 min in PBST and then mounted with Vectashield. For detection of GFP::MSH-5, worms were dissected and fixed in 1× EGG buffer containing 0.1% Tween (instead of PBST). Detection of [RPA-1::YFP] was performed as previously described (85). Primary antibodies used in this study were: mouse monoclonal anti-HA tag (pre-absorbed on N2 worms to reduce non-specific binding; 1:1000 dilution; Covance), rabbit anti-HA tag (1:250 dilution; Invitrogen), rabbit anti-BRD-1 (1:500 dilution) (50), chicken anti-SYP-1 (1:400 dilution) (51), guinea pig anti-HTP-3 (1:500 dilution) (60), mouse monoclonal anti-GFP (1:500 dilution; Roche), guinea pig anti-ZHP-3 (1:500 dilution) (21), rabbit anti-OLLAS tag (pre-absorbed on N2 worms to reduce non-specific binding; 1:1500 dilution; GenScript), rabbit anti-RAD-51 (1:10,000 dilution; SDIX) and rabbit anti-PLK-2 (1:500 dilution) (86). Appropriate secondary antibodies were conjugated with Alexa Fluor 488 or 594 (1:500 dilution) or with Alexa Fluor 647 (1:250 dilution). Images were collected as z-stacks (0.3 μm intervals) using an UPlanSApo 100x NA 1.40 objective on a DeltaVision System equipped with a CoolSNAP HQ2 camera. Files were deconvolved with SoftWORx software and processed in Adobe Photoshop, where some false colouring was applied. Samples acquired by super-resolution microscopy (Fig 3C) were prepared as previously reported (63) without modifications and imaged with a DeltaVision OMX.

### Biochemistry

For whole-cell protein extraction, 200 age-matched animals (24 hours post-L4 stage) were picked into 1× Tris-EDTA buffer (10 mM Tris pH 8, 1 mM EDTA) containing 1× protein inhibitor cocktail (Roche), snap-frozen in liquid nitrogen. After thawing, an equal volume of 2× Laemmli buffer was added. Samples were boiled for 10 min, clarified and separated on pre-cast 4–20% gradient acrylamide gels (Bio Rad).

Fractionated protein extracts for western blotting and immunoprecipitation were prepared as previously reported (51). Western blotting used 50 μg protein samples from each fraction, whereas immunoprecipitation assays used at least 1 mg samples of pooled soluble nuclear and chromatin-bound fractions. Proteins were transferred onto nitrocellulose membrane for 1 hour at 4°C at 100V in 1× Tris-glycine buffer containing 20% methanol. Membranes were blocked for 1 hour in 1× TBS containing 0.1% Tween (TBST) and 5% milk; primary antibodies were added into the same buffer and incubated overnight at 4°C. Membranes were then washed in 1× TBST and then incubated with appropriate secondary antibodies in TBST containing 5% milk for 2 hours at room temperature. After washing, membranes were incubated with ECL and developed with a ChemiDoc system (BioRad). To detect phosphorylated CHK-1^S345^, TBST containing 5% BSA (instead of milk) was used for blocking and antibody dilution. The following antibodies were used for western blotting: mouse monoclonal anti-HA tag (1:1000 dilution; Cell Signalling), rabbit anti-HA tag (1:500 dilution; Invitrogen), chicken anti-GFP (1:4000 dilution; Abcam), mouse anti-GAPDH (1:5000 dilution; Ambion), rabbit anti-Histone H3 (1:100,000 dilution; Abcam); rabbit anti-phospho-CHK-1^S345^ (1:1000 dilution; Cell Signalling), HRP-conjugated anti-mouse (1:2500 dilution) and anti-rabbit (1:25,000 dilution; both Jackson ImmunoResearch) and HRP-conjugated anti-chicken (1:10,000 dilution; Santa Cruz).

### Irradiation

Age-matched worms (24 hours post-L4 stage) were exposed to the indicated dose of IR with a Gammacell irradiator containing a ^137^Cs source. For viability screening, irradiated worms were allowed to lay eggs for 24 hours and then removed; hatched versus unhatched eggs were scored the following day. For cytological analysis, worms were dissected and immunostained at the indicated times.

### CRISPR-Cas9 Tagging

All the details relative to the tagging strategy followed to generate the *brc-1::HA, OLLAS::cosa-1* and *GFP::msh-5* lines are available upon request.

## Acknowledgements

We are grateful to S. Boulton, N. Bhalla, E. Martinez-Perez, M. Zetka and R. Lin for kindly providing the anti-BRD-1, anti-ZHP-3, anti-SYP-1, anti-HTP-3 and anti-PLK-2 antibodies respectively. We thank A. Graf for performing the microinjections, and J. Yanowitz and J. Engebrecht for sharing unpublished data. Some strains were provided by the CGC, which is funded by the NIH Office of Research Infrastructure Programs (P40 OD010440). NS is the recipient of an Interdisciplinary Cancer Research fellowship (INDICAR) funded by the Mahlke-Obermann Stiftung and the European Union’s Seventh Framework Programme for Research, Technological Development under grant agreement no 609431. Research in VJ laboratory is funded by the Austrian Science Fund (project no. SFB F3415-B19). MRDS was supported by the Austrian Science Fund doctoral programme (project no. W1238).

## Author contributions

Conceptualization: NS.; Funding acquisition: NS.; Investigation: NS, EJ, MRDS.; Methodology: NS.; Project administration: NS.; Resources: NS VJ.; Visualization: NS. Writing (original draft): NS.

## Supplementary figure captions

**S1 Fig.**
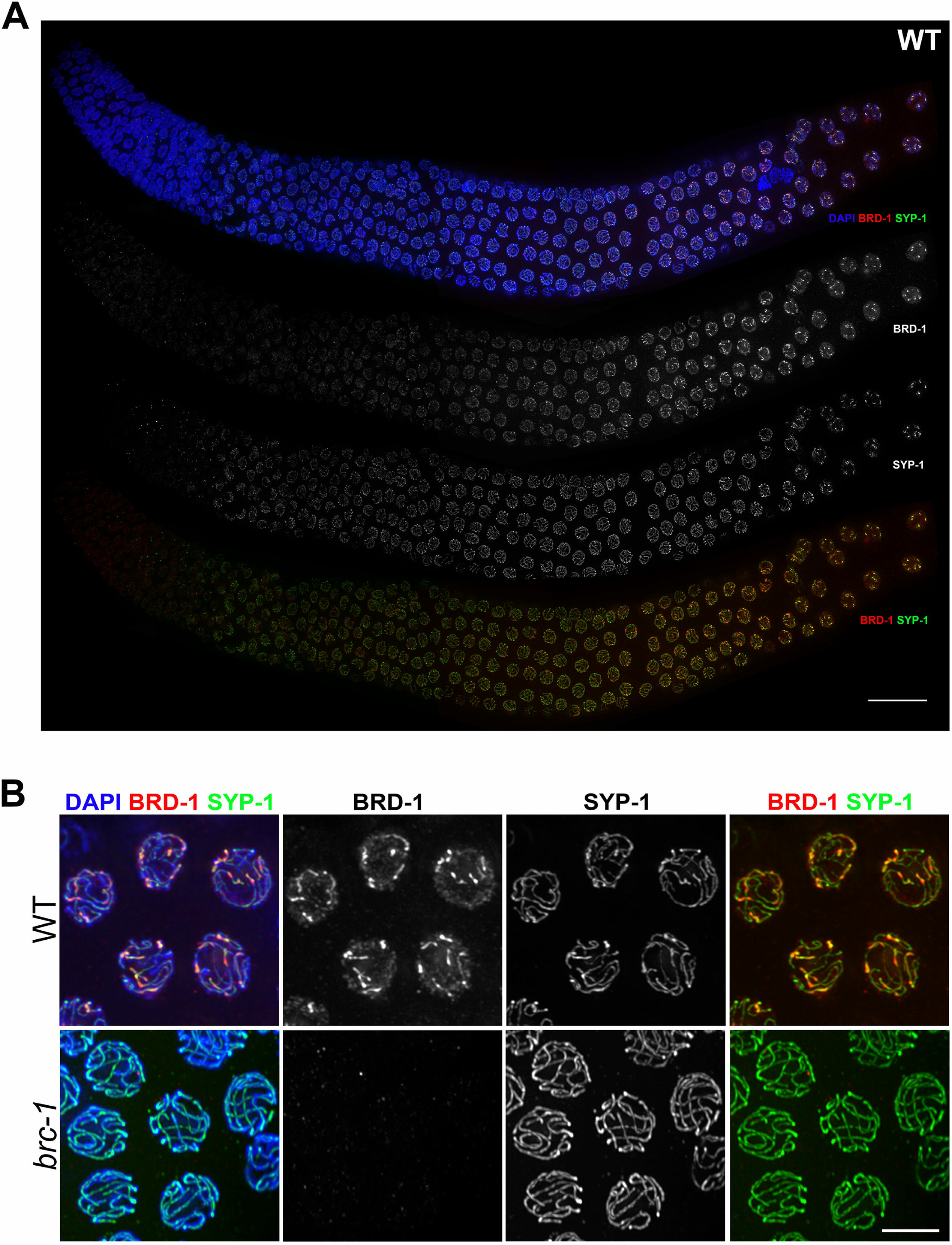
BRD-1 and BRC-1 localization patterns are identical during meiotic prophase I. (A) BRD-1 and SYP-1 immunostaining in wild-type animals shows that the BRD-1 expression pattern is identical to the one observed for BRC-1::HA. Note enrichment on the SC and retraction to the short arms of bivalents. Scale bar, 30 μm. (B) Interdependent DNA loading for BRD-1 and BRC-1 is shown by a lack of DNA-binding by BRD-1 in *brc-1* mutant germlines. Scale bar, 5 μm.

**S2 Fig.**
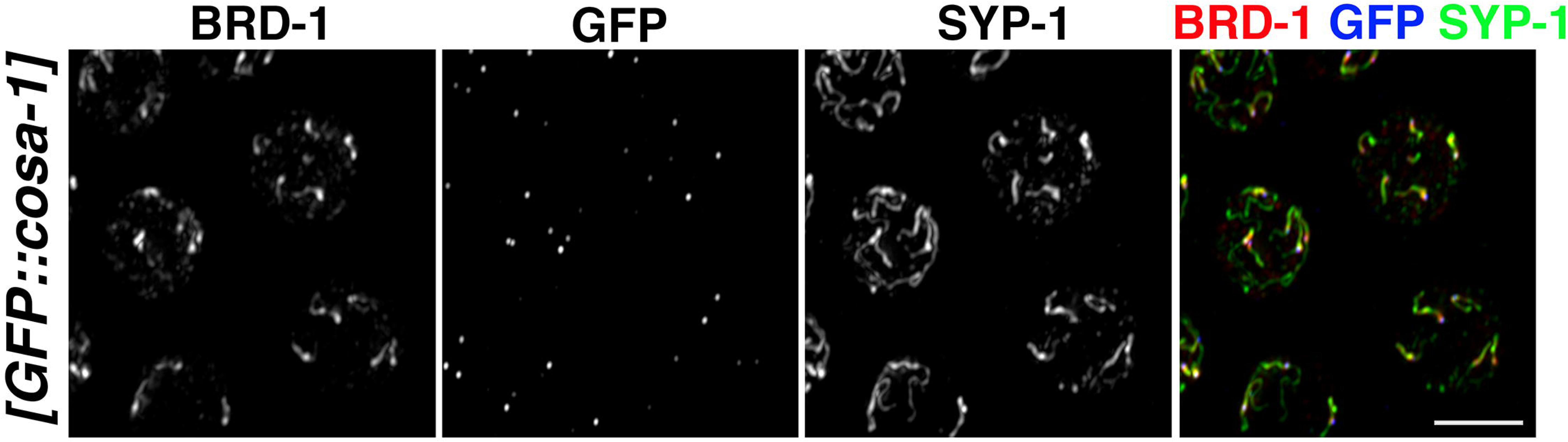
BRD-1 is enriched at chromosome subdomains containing presumptive CO sites. Late pachytene nuclei of *[GFP*::*cosa-1]* animals were stained for BRD-1, GFP and SYP-1. As previously observed for BRC-1, BRD-1 is progressively enriched at regions surrounding the CO site. Scale bar, 5 μm.

**S3 Fig.**
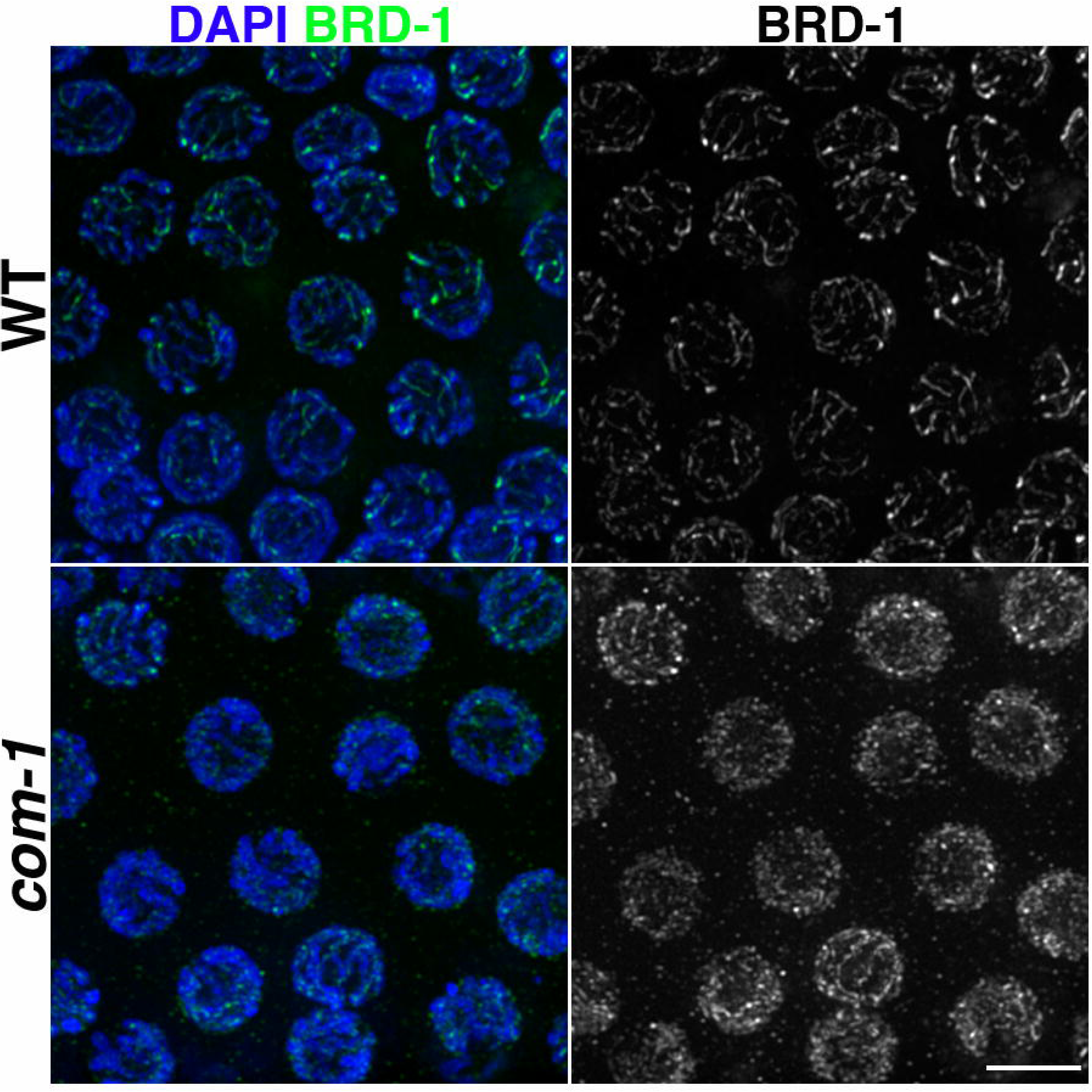
Association of BRD-1 with the SC is largely disrupted in DSBs resection-defective *com-1* mutants. Mid-/late pachytene nuclei of the wild type (WT) and *com-1* mutant were stained for BRD-1. BRD-1 loading onto the SC is drastically reduced when DNA resection is impaired. Scale bar, 5 μm.

**S4 Fig.**
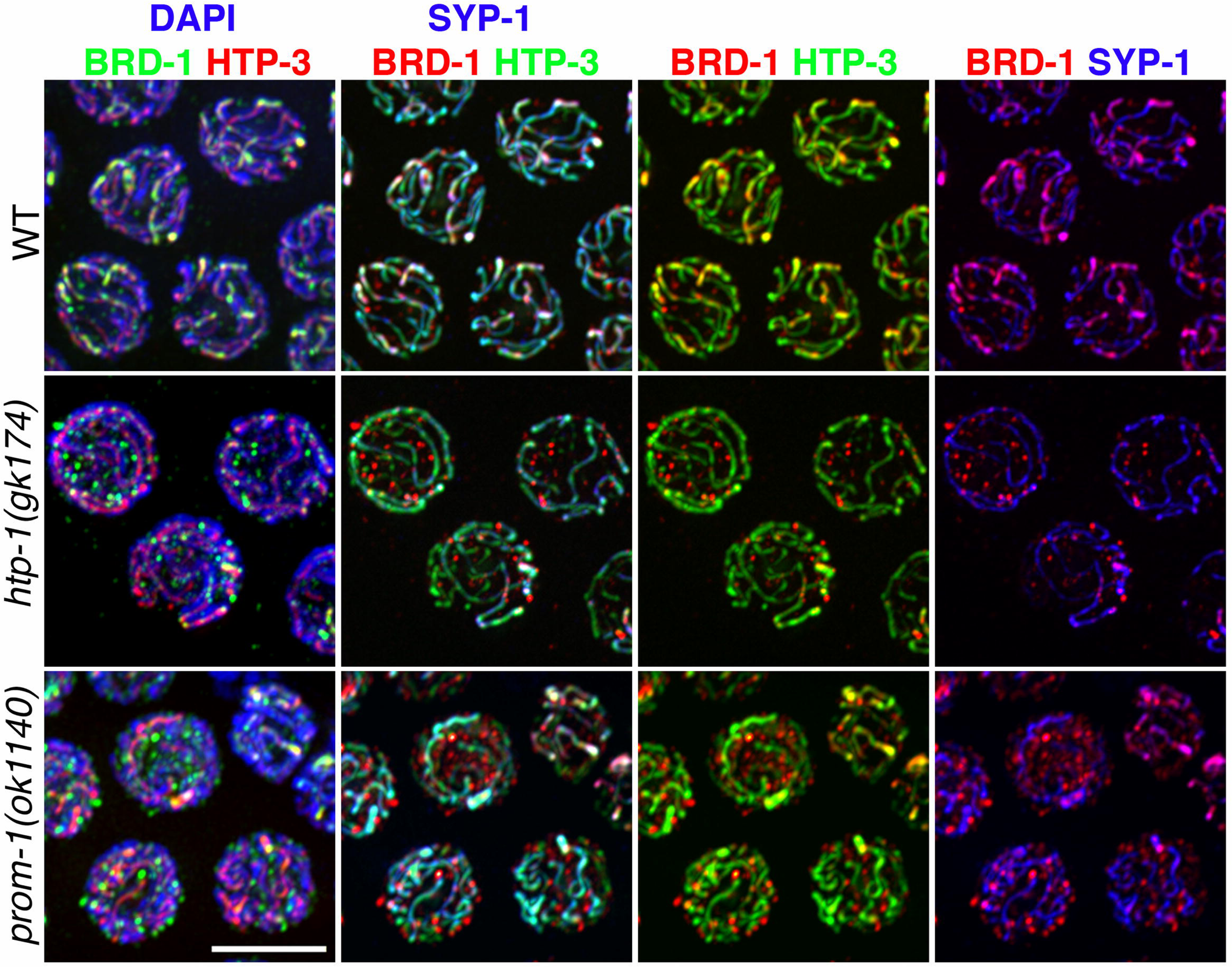
Non-homologous synapsis largely impairs loading of BRD-1, leading to nucleoplasmic accumulation. Late pachytene nuclei in the wild-type (WT) and *htp-1* and *prom-1* mutants were stained with BRD-1, SYP-1 and HTP-3. In both mutants, BRD-1 is largely excluded from the SC and forms nucleoplasmic agglomerates. Scale bar, 5 μm.

**S5 Fig.**
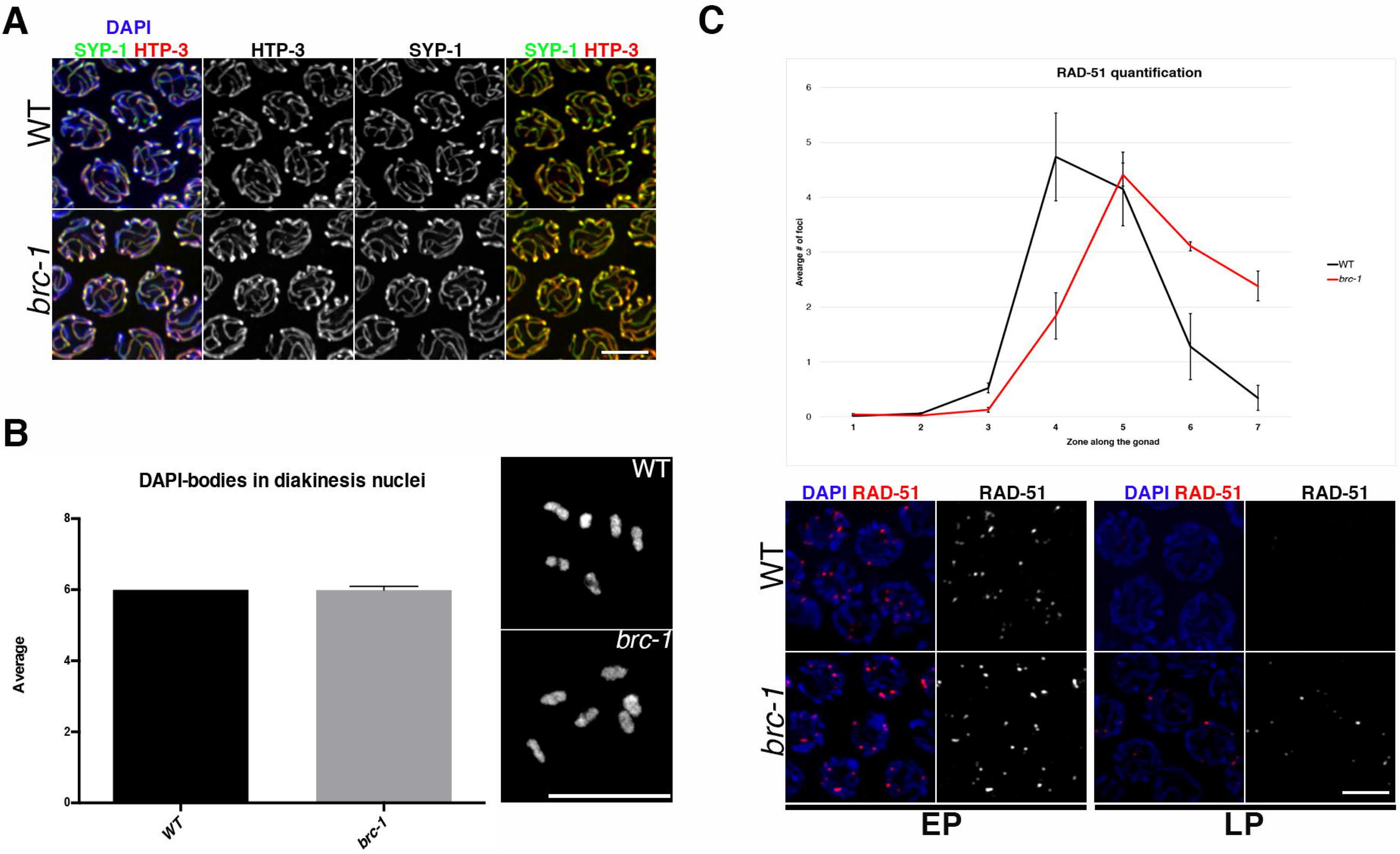
Synapsis and chiasmata formation occur normally but RAD-51 foci accumulate in the *brc-1* mutant. (A) SC assembly in the wild type (WT) and *brc-1* mutant was monitored by co-staining for the axial element HTP-3 and central element SYP-1. The *brc-1* mutant had no obvious defect in establishing synapsis. Scale bar, 5 μm. (B) DAPI-staining of diakinesis nuclei does not show defective chiasmata formation in the *brc-1* mutant. Scale bar, 5 μm. Number of diakinesis nuclei analysed: WT, 44; *brc-1* mutant, 46. Error bars show standard deviation. (C) Top: quantification revealed an accumulation and delayed disappearance of RAD-51 foci in the absence of BRC-1. Bottom: representative nuclei from early pachytene (EP) and late pachytene (LP) stages stained with DAPI and anti-RAD-51 antibody. Scale bar, 5 μm. For RAD-51 foci quantification, the following numbers of nuclei were counted in each region in WT (and the *brc-1* mutant): zone 1, 154 (173); zone 2, 226 (189); zone 3, 189 (186); zone 4, 157 (156); zone 5, 113 (136); zone 6, 95 (125); zone 7, 92 (72). Error bars show S.E.M.

**S6 Fig.**
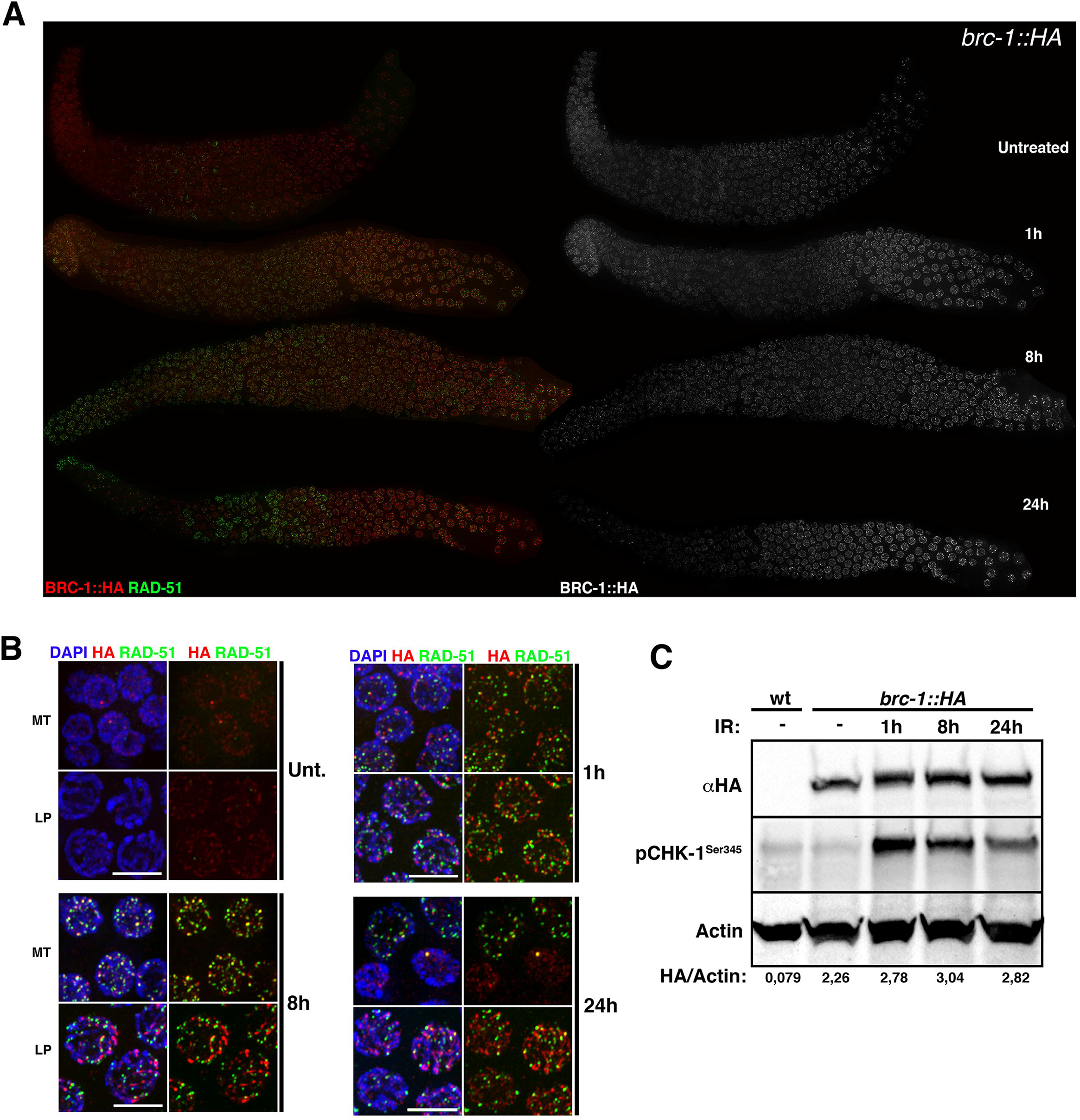
Exogenous DNA damage increases BRC-1 levels and triggers its association with RAD-51 in mitotic nuclei. (A) Whole-mount gonads of irradiated and non-irradiated *brc-1*::*HA* worms immunostained for HA and RAD-51. Animals were exposed 75 Gy IR and analysed at the indicated time points. (B) Representative nuclei from the pre-meiotic region (MT) and late pachytene (LP) stage of gonads analysed at different times after IR. Note BRC-1::HA focus formation in pre-meiotic nuclei, along with robust co-localization with RAD-51 at 8 hours and occasionally at 24 hours post irradiation. Scale bars, 5 μm. (C) Western blot analysis of whole-cell extracts show a shift in BRC-1::HA migration after irradiation. Wild-type (wt) worms were the negative control. Actin was the loading control and induction of phosphorylated CHK-1^Ser345^ was used as a positive control for irradiation. The ratio of BRC-1::HA to actin (HA/Actin) is shown as an abundance index.

